# Progressive Centro-Parietal ERP Responses During Contextual Integration Across Symbolic Domains

**DOI:** 10.64898/2026.04.25.720770

**Authors:** María Guadalupe Yáñez-Ramos, Daniel Zarabozo Enríquez de Rivera, Andrés Antonio González Garrido

## Abstract

Many cognitive processes depend on integrating information as it becomes available to construct meaningful interpretations. Prior work has shown graded and incremental context effects, especially in language, but it remains less clear whether contextual integration exhibits a comparable temporal profile across symbolic domains when structured input is examined within congruent sequences. Twenty-seven participants processed congruent four-element sequences designed to be structurally comparable across lexical, algebraic, and graphical domains while event-related potentials were recorded.

In the 250–500 ms interval, mean amplitudes increased systematically with sequence position within a predefined centro-parietal region of interest (p < .001). The Domain × Position interaction did not reach significance (p = .056), although modest domain-related differences in the buildup profile cannot be ruled out. A follow-up analysis showed that the increase to the response-relevant final position was larger than earlier increases (p < .001). Additional analyses indicated maximal amplitudes over parietal sites and the clearest graded increase over central sites.

These findings indicate that context-sensitive activity was progressive but not uniform across sequence positions, with the strongest increase occurring when the sequence reached its final, response-relevant completion point. The presence of position-related increases across lexical, algebraic, and graphical domains is consistent with the view that centro-parietal ERP activity in the 250–500 ms window tracks the progressive buildup of contextual integration during structured sequence processing.

**Highlights:**

- Context-sensitive ERP activity increased across sequence position.
- The strongest increase occurred at the final completion point.
- Maximal amplitudes were observed over parietal electrodes.
- Central sites best captured graded position-related modulation.
- Position-related buildup was observed across symbolic domains.

## Introduction

Constructing meaningful interpretations requires integrating contextual information, a process that plays a central role across many cognitive domains, including language comprehension, mathematical reasoning, and graphical pattern processing. A large body of electrophysiological literature has shown that neural responses in the 250–500 ms latency range are sensitive to contextual information during comprehension (Kutas & Hillyard, 1980; Kutas & Federmeier, 2011). More recent accounts have emphasized that comprehension unfolds continuously, with incoming input evaluated relative to an evolving internal representation rather than only at isolated target positions (Federmeier, 2022).

In this time range, effects are shaped not only by incongruity but also by graded forms of contextual support and lexical prediction (Van Petten & Luka, 2012; Brothers et al., 2015). Predictability and incongruity do not necessarily produce identical effects in this interval (Lau et al., 2016), and responses vary with graded aspects of contextual support, including predictability and contextual constraint (Van Petten & Luka, 2012; Federmeier, 2022; Dambacher et al., 2006). Classic sentence-processing studies showed that word position within a sentence is associated with systematic changes in N400 amplitude, and that the influence of lexical variables, such as word frequency, is reduced as contextual constraints increase (Van Petten & Kutas, 1990, 1991). More recent work using item-level and mixed-effects approaches has reinforced the view that processing changes continuously as sequential input unfolds (Payne et al., 2015; Payne & Federmeier, 2018). Computational accounts are broadly consistent with this perspective, modeling activity in this time range as gradual updating during the accumulation of contextual information (Rabovsky et al., 2018; Michaelov et al., 2024; Nour Eddine et al., 2024). Taken together, these findings suggest that context effects can build up incrementally, especially in language.

Although this literature is most developed in language, related findings extend beyond the verbal domain. N400-like effects have been reported for arithmetic expressions, suggesting sensitivity to knowledge-based or meaning-related relations during mathematical processing (Niedeggen et al., 1999; Dickson et al., 2018). Research on visual narratives has likewise shown that comprehension of sequential images depends on integrating information across panels over time, and ERP (event-related potential) studies in that domain have examined how sequential structure and ordinal position shape online processing (Cohn et al., 2012, 2014). This broader perspective is also supported by evidence that visual context can incrementally influence comprehension and can be tracked with ERPs, even in paradigms that involve explicit verification responses (Knoeferle et al., 2011). Together, these findings raise the possibility that context-sensitive activity during structured sequence processing is not exclusive to language.

What remains less clear is whether, when structured input is examined within congruent sequences, contextual integration exhibits a comparable temporal profile across symbolic domains, and whether such a profile may reflect a common electrophysiological pattern associated with constructing meaningful interpretations. Most prior work has relied on contrasts between expected and unexpected inputs, whereas the present question concerns how neural responses develop as contextual support is preserved and accumulates across successive elements. This distinction matters because it allows us to ask not only whether context effects are present, but also how they develop across the sequence, whether this development is uniform, and whether a similar temporal profile is observed across domains. We therefore focused on congruent sequences to examine the buildup of contextual processing without making expectancy violation the primary source of the effect.

Although the 250–500 ms interval overlaps with the canonical N400 time range, the present analysis was not designed to estimate a traditional N400 incongruity effect. Classic N400 effects are typically defined as amplitude differences between incongruent and congruent, or between less predictable and more predictable stimuli. In contrast, the present study focused specifically on congruent sequences because these are the trials in which incoming information can be progressively integrated into a coherent interpretation. We therefore avoid referring to the observed modulation as a conventional N400 effect and instead describe it as context-sensitive ERP activity in the 250–500 ms time window.

The present study addressed this question using congruent four-element sequences in three visual domains: lexical sentences, algebraic operations, and graphical compositions. By using rigorously controlled four-element sequences with the same bounded structure across domains, the design enabled comparisons of contextual buildup while minimizing differences in sequence length and overall format. We asked whether responses in the 250–500 ms interval would increase across sequence positions, and whether this increase would follow a comparable profile across domains. Based on prior work on graded context effects, we predicted progressive increases from early to late sequence positions. At the same time, we asked whether this progression would be uniform or whether the strongest increase would emerge at sequence completion. Such a pattern would suggest that contextual integration over structured input is progressive but not uniform. It would also help constrain accounts of sequence comprehension beyond categorical target responses and raise the possibility that similar electrophysiological dynamics support contextual integration across symbolic domains.

## Method

### Participants

Thirty undergraduate students participated in the experiment. Data from three participants were excluded (one incomplete recording and two excessive movement artifacts), leaving a final sample of 27 participants (21 male, 6 female; M = 21.0 years, SD = 1.9). All participants were native Spanish speakers, had normal or corrected-to-normal vision, were right-handed as assessed with Annett’s questionnaire (Annett, 1970), and reported no history of neurological or psychiatric disorders. Participants had been previously assessed to ensure that they had the necessary knowledge to solve algebraic operations with monomials. All participants provided written informed consent. The study was approved by the Ethics Committee of the Institute of Neuroscience, University of Guadalajara.

### Stimuli

The task was programmed and presented in PsychoPy (Peirce, 2009). Three types of four-element sequences were presented visually: lexical, algebraic, and graphical. Across domains, sequences were designed to be as structurally comparable as possible, with the first three elements establishing a progressively constraining context and the fourth serving as the critical completion item. Elements A, B, and C were presented in white (Figure 1), whereas element D was presented in yellow to indicate that it was the final element of the sequence and the point at which participants made their behavioral response. On 50% of trials, the fourth element was congruent with the preceding context; on the remaining trials, it was incongruent. A total of 240 trials were presented.

**Figure 1.**
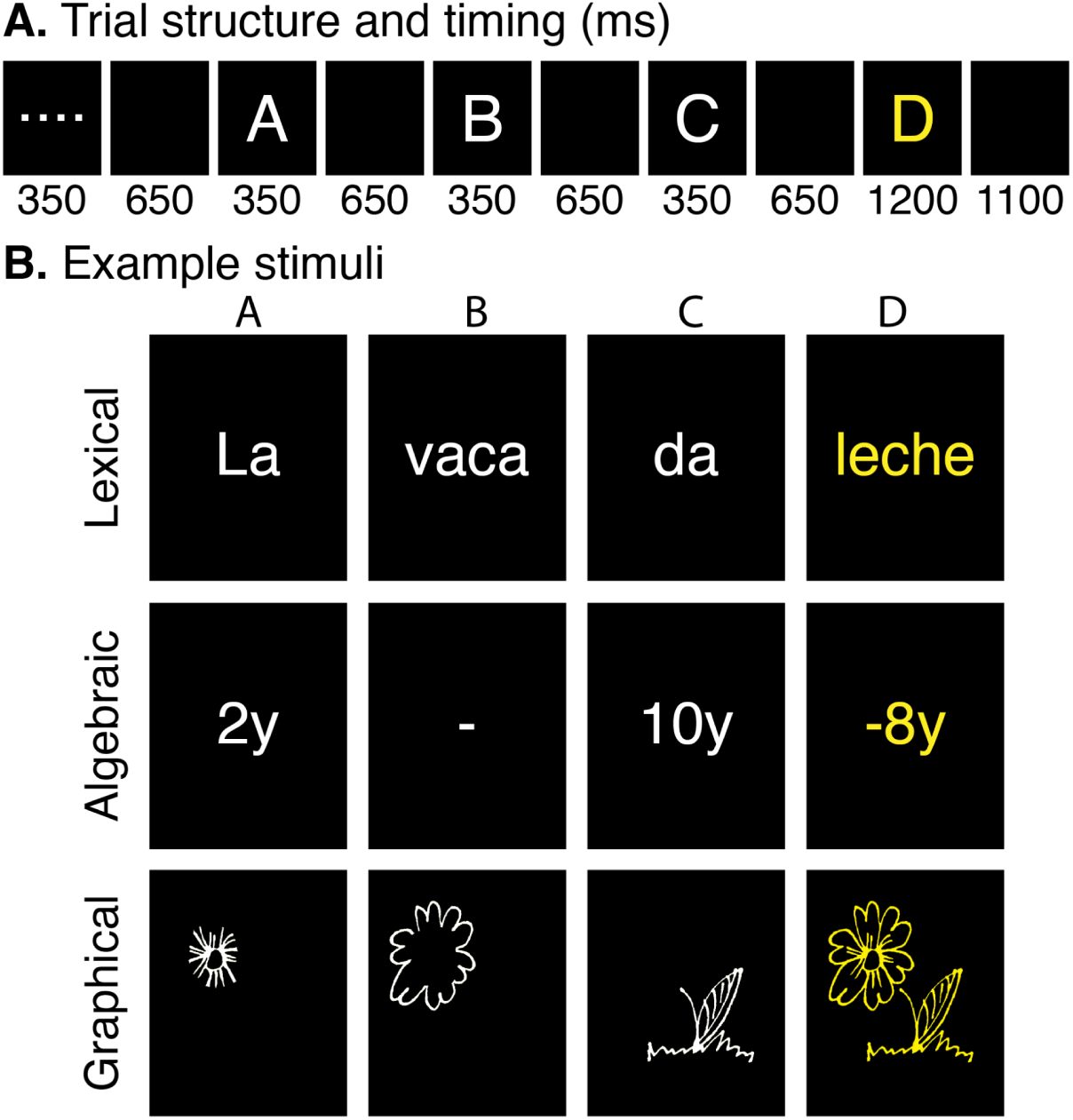
Trial Structure and Example Stimuli. (A) Temporal structure of a single trial. Durations are shown in milliseconds. Each trial began with a 350 ms fixation display (four white dots), followed by sequential presentation of elements A–C, each displayed for 350 ms and separated by 650 ms interstimulus intervals. Finally, the fourth element was presented for 1200 ms. Each trial ended with a 1100 ms blank screen. (B) Example sequences from the three stimulus domains (lexical, algebraic, and graphical). The fourth element was displayed in yellow during the experiment to indicate the response window. The lexical example is shown in Spanish, the language used for stimulus presentation (“La vaca da leche”; “The cow gives milk”).

The lexical stimuli consisted of four-word sentences derived from a published stimulus set (Yáñez-Ramos et al., 2022), in which the final word could be congruent or incongruent with the preceding context. The mean cloze probability for congruent lexical stimuli was .86. This high cloze value was selected to make lexical completions as constrained as possible, given that algebraic and graphical sequences were designed to have determinate correct completions.

The algebraic stimuli consisted of operations with monomials, including a first monomial, an operator, a second monomial, and a proposed solution that could be correct or incorrect. The graphical stimuli consisted of line-based drawings in which the first three elements were partial components of an image and the fourth element was either the integrated image corresponding to those components or an unrelated image. Stimuli were presented centrally on a 22-inch monitor (1680 × 1050 resolution) at an approximate viewing distance of 60 cm. Lexical and algebraic stimuli were presented as PsychoPy TextStim objects in normalized screen units and centered at fixation, whereas graphical stimuli were presented as images (600 × 600 pixels, scaled to 200 × 200 pixels), subtending approximately 5.4° × 5.4° of visual angle.

To ensure task engagement, participants judged whether the fourth element was congruent or incongruent with the preceding elements by pressing the left mouse button for congruent responses and the right mouse button for incongruent responses. The fourth element was presented in yellow to indicate that it was the final element in the sequence and the point at which the response was required.

### Procedure

Participants were seated at a viewing distance of 60 cm from a computer screen. The session began with a practice phase comprising 15 trials (5 per stimulus type) to familiarize participants with the task. The experimental session comprised 240 trials distributed across three stimulus domains: Lexical, Algebraic, and Graphical. Each domain contributed 80 trials, divided into four blocks of 20 trials, allowing participants to take short breaks between blocks to minimize fatigue. The three stimulus domains were presented in counterbalanced order across participants to control for potential carryover effects. Within each block, trials were pseudorandomized such that no more than three congruent or incongruent trials occurred consecutively.

Each trial began with a fixation display consisting of four white dots presented for 350 ms, followed by a 650 ms interstimulus interval (ISI). The four sequence elements (A–D) were then presented sequentially. Elements A, B, and C were displayed for 350 ms each in white and separated by 650 ms ISIs. The fourth element was presented for 1200 ms in yellow. The color change and longer presentation duration were used to indicate the response point clearly and to allow sufficient time for participants to evaluate the completion item while it remained visible. Participants responded as soon as they determined whether it was congruent or incongruent with the preceding elements. Each trial ended with a 1100 ms blank screen. The total duration of each trial was approximately 6.3 s (Figure 1).

### EEG Recording and Data Analysis

The electroencephalogram (EEG) was recorded using a Neuronic Medicid IV system (Neuronic Mexico) from scalp electrodes placed at frontal (F3, Fz, F4), central (C3, Cz, C4), parietal (P3, Pz, P4), and occipital (O1, O2) sites according to the International 10-20 System. A biauricular reference was used with electrodes placed on the left and right earlobes and electrically bridged. The ground electrode was placed on the forehead. Horizontal and vertical eye movements were monitored using bipolar electrooculogram (EOG) recordings, with one electrode placed lateral to the right eye’s outer canthus and another below the left eye. Signals were recorded in single-ended mode and digitized at a sampling rate of 200 Hz. The amplifier gain was set to 10,000, with an online band-pass filter of 0.1-30 Hz and a 60-Hz notch filter applied to reduce line noise. Electrode impedances remained below 10 kΩ throughout the recording.

### ERP Preprocessing

Continuous EEG data were imported from the Neuronic system into EEGLAB (Delorme & Makeig, 2004) for preprocessing. Offline filtering was applied using a zero-phase Butterworth band-pass filter (0.1-18 Hz, 2nd order), following recommendations for ERP research (Tanner et al., 2015; Luck, 2014). ERP analyses were conducted using ERPLAB (Lopez-Calderon & Luck, 2014). Data were segmented into epochs from −100 to 500 ms relative to stimulus onset and baseline-corrected using the prestimulus interval.

Trials with incorrect responses, omissions, or ocular artifacts were excluded from analysis. Artifact rejection was performed in EEGLAB/ERPLAB using automated procedures. Ocular artifacts were identified using a cross-covariance-based blink detection algorithm, and additional moving-window criteria were applied to detect abrupt voltage steps, excessive peak-to-peak activity, and flatline segments. Trials flagged by any artifact or behavioral exclusion were removed prior to ERP averaging. Across subjects and sequence elements, the mean proportion of rejected epochs was 23.3%. Rejection rates ranged from 21.1% in the algebraic domain to 24.9% in the lexical domain.

After artifact rejection, the number of retained trials was comparable across domains and sequence positions, with means ranging from 29 to 32 trials per domain and element, and no systematic differences across conditions. For each participant, averaged ERP waveforms were computed separately for each sequence element (A, B, C, and D) within each stimulus domain. Only congruent trials were included in the present ERP analyses in order to examine the buildup of contextual processing within interpretable sequences.

### Statistical Analyses

#### Behavioral Analysis

Behavioral performance was analyzed to confirm task engagement and to assess potential differences in task demands across domains. Accuracy (percentage of correct responses) and reaction times (RTs) for correct responses were computed separately for each domain across the full task. Reaction times were measured from the onset of the fourth element (D), when participants indicated whether the sequence was congruent or incongruent. Trials with omissions or reaction times below 300 ms were excluded as anticipatory responses (0.7% of trials), as such latencies are unlikely to reflect stimulus-driven decision processes in this task. Accuracy was calculated on the remaining trials, and RT analyses included only correct responses.

Repeated-measures analyses of variance (ANOVAs) with Domain (lexical, algebraic, and graphical) as a within-subject factor were conducted separately for accuracy and RTs using IBM SPSS Statistics (Version 25). Sphericity was evaluated using Mauchly’s test, and Greenhouse-Geisser corrections were applied when necessary. Effect sizes are reported as partial eta squared (ηp²), and Bonferroni-adjusted pairwise comparisons were used for post hoc tests.

#### ERP Statistical Analysis

ERP mean amplitudes were analyzed in two predefined time windows corresponding to early sensory processing (0–180 ms) and later context-sensitive processing (250–500 ms). Statistical analyses were conducted using linear mixed-effects models implemented in SPSS with restricted maximum likelihood (REML) estimation. Fixed effects were evaluated using Type III *F*-tests.

##### Early sensory analysis (element A: 0–180 ms)

Mean amplitude was quantified within a 0–180 ms time window. This window was selected a priori to characterize early sensory-perceptual processing, as visual ERP activity in this range is commonly associated with early components such as the P1 and N1 and with sensitivity to low-level stimulus properties (Rokszin et al., 2016), rather than with later context-sensitive effects (Kutas & Federmeier, 2011). To characterize early topographic response patterns, electrodes were grouped into four anterior-posterior regions: frontal (F3, Fz, F4), central (C3, Cz, C4), parietal (P3, Pz, P4), and occipital (O1, O2). Mean amplitudes were computed for each region and entered into linear mixed-effects models with fixed effects of Domain (three levels), Region (four levels), and their interaction. A subject-level random intercept was included to account for repeated measurements within participants. In an additional analysis, Region was modeled as an ordinal predictor to test for a linear posterior-to-anterior gradient across scalp regions.

##### Contextual integration analysis (250–500 ms)

Mean amplitudes were quantified within a 250–500 ms time window, selected a priori based on prior work showing robust context-sensitive effects in this latency range during comprehension, including graded effects of predictability and contextual constraint. Analyses focused on a predefined centro-parietal region of interest (ROI) comprising electrodes C3, Cz, C4, P3, Pz, and P4, motivated by prior literature indicating that context-sensitive activity in this time range typically exhibits a centro-parietal distribution (Kutas & Federmeier, 2011; Van Petten & Luka, 2012). For the primary analysis, mean amplitudes from the centro-parietal ROI were entered into linear mixed-effects models with fixed effects of Sequence Position (A–D), Domain (Algebraic, Graphical, Lexical), and their interaction. Sequence Position was modeled as an ordinal predictor (coded 0-3), corresponding to increasing contextual information across the sequence. A random slope for Position by subject was included to account for inter-individual variability in the position effect.

Because the primary model tested whether responses increased across sequence positions, a follow-up analysis was conducted to determine whether the increase was uniform or disproportionately larger at the final sequence step. For each participant and domain, adjacent difference scores were computed within the centro-parietal ROI (B–A, C–B, and D–C). These transition scores were entered into a linear mixed-effects model with Domain and Transition as fixed effects and subject as a random intercept. Bonferroni-corrected pairwise comparisons were used to test whether the increase at the final transition (D–C) differed from earlier transitions.

To further characterize the spatial distribution of the position effect, an additional mixed-effects analysis included region of interest (ROI: frontal, central, parietal, occipital) as a factor. In this model, mean amplitudes were entered into a linear mixed-effects model with fixed effects of Domain, ROI, Sequence Position, and their interactions, while retaining a random slope for Position by subject. This analysis was intended to assess whether the strength of the position effect varied across scalp regions and to determine whether regions showing maximal response amplitude also exhibited the strongest position-related modulation. Transition-level follow-up analyses were restricted to the predefined centro-parietal ROI used in the main late-window analysis.

For visualization, Figure 3A and 3C display amplitudes expressed relative to position A within each participant and domain (A = 0), whereas the scalp topographies in Figure 3B were computed from the absolute mean amplitude elicited by element D in a 50-ms window centered on the grand-average peak (∼350 ms; 325–375 ms).

## Results

### Behavioral Performance

Mean accuracy was 98.00% (*SD* = 1.95%) in the lexical domain, 96.98% (*SD* = 3.20%) in the algebraic domain, and 97.71% (*SD* = 2.50%) in the graphical domain. A repeated-measures ANOVA revealed no significant effect of Domain on accuracy, *F*(2, 52) = 1.18, *p* = .317, ηp² = .043.

Mean reaction times (RTs) for correct trials were 889 ms (*SD* = 110 ms) in the lexical domain, 828 ms (*SD* = 130 ms) in the algebraic domain, and 805 ms (*SD* = 110 ms) in the graphical domain. A repeated-measures ANOVA showed a significant main effect of Domain, *F*(2, 52) = 17.93, *p* < .001, ηp² = .408. Bonferroni-adjusted comparisons indicated that responses were slower in the lexical domain than in the algebraic domain (mean difference = 61 ms, *p* = .001) and the graphical domain (mean difference = 84 ms, *p* < .001), whereas the algebraic and graphical domains did not differ, *p* = .493.

According to predefined exclusion criteria, 0.7% of trials were removed due to omissions or anticipatory responses (<300 ms). Among the remaining trials, incorrect responses accounted for 2.4%.

### ERP Results

Grand-average ERP waveforms for each sequence element (A–D) across scalp regions are shown in Figure 2. Early activity (0–180 ms) was maximal over occipital sites and decreased anteriorly. In contrast, activity in the 250–500 ms window showed a centro-parietal maximum that increased progressively across sequence elements, peaking at element D in all three domains. Maximal positivity occurred at approximately 350 ms over parietal electrodes.

**Figure 2.**
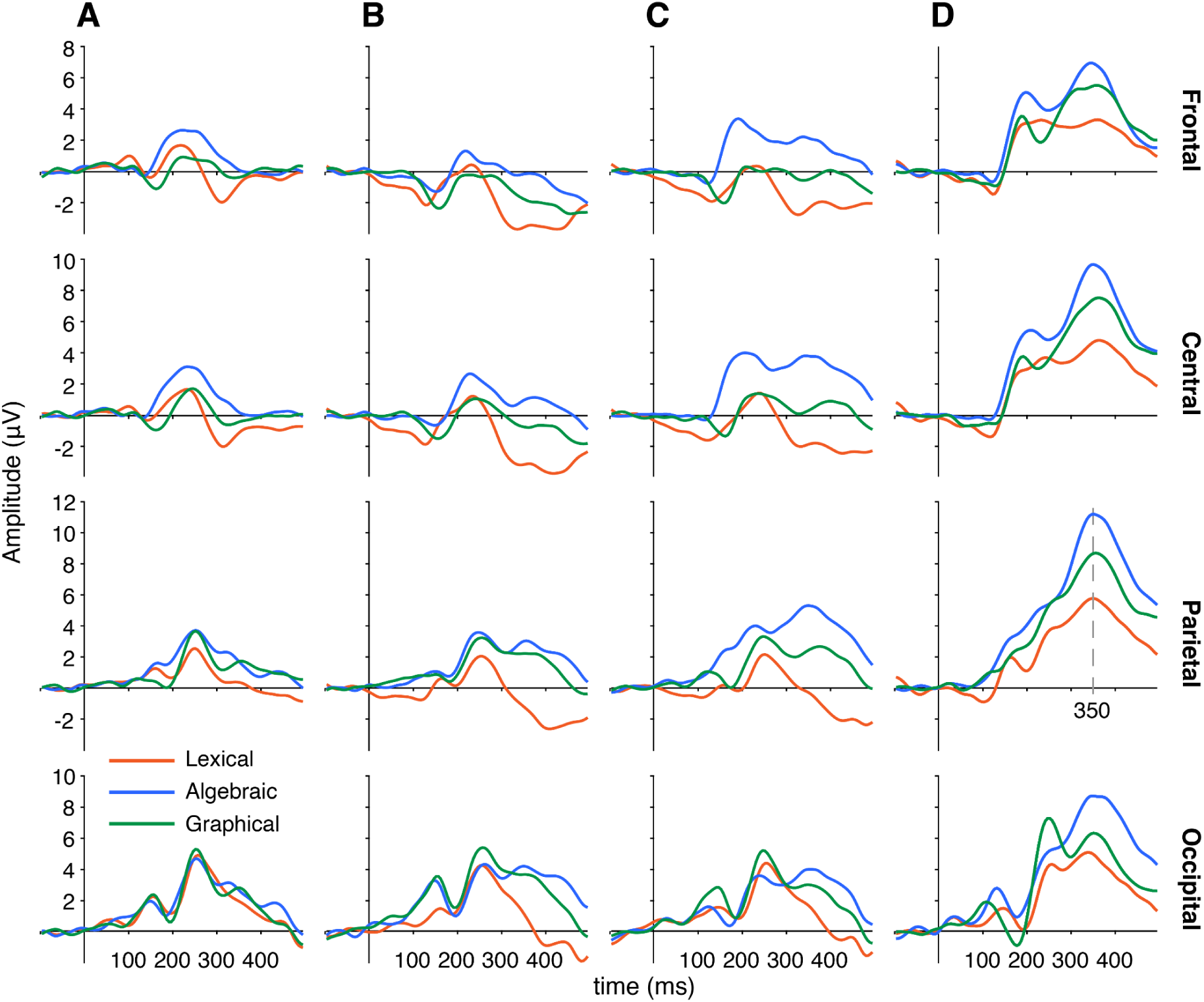
Grand-Average ERPs Across Sequence Elements and Scalp Regions. Grand-average ERP waveforms elicited by each sequence element (A–D; columns) across scalp regions (rows: frontal, central, parietal, and occipital). Waveforms represent the average of predefined electrode clusters within each region (see Methods). Red = lexical sequences, blue = algebraic, green = graphical. ERPs are time-locked to stimulus onset (0 ms) and baseline-corrected to the prestimulus interval. Amplitude is plotted in microvolts (µV) and time in milliseconds (ms). The gray dashed line in the parietal D panel indicates the approximate latency of the maximal late positivity (∼350 ms), corresponding to the peak within the 250–500 ms analysis window.

#### Early Sensory Response to Element A (0–180 ms)

A linear mixed-effects model with Region modeled as a categorical factor revealed a significant main effect of Region, *F*(3, 286) = 9.89, *p* < .001, indicating differences in amplitude across scalp regions. No Domain × Region interaction was observed, *F*(6, 286) = 0.36, *p* = .905, suggesting that the early spatial distribution did not differ across domains.

In a secondary analysis, Region was modeled as an ordinal predictor, revealing a significant posterior-to-anterior gradient, *F*(1, 292) = 24.73, *p* < .001. Amplitudes decreased systematically from occipital to frontal regions (β = −0.26 µV per region step, 95% CI [−0.45, −0.06]). This gradient did not interact with Domain, *F*(2, 292) = 0.82, *p* = .442, indicating a consistent early topographic distribution across domains.

#### Late-Window Centro-Parietal Effects (250–500 ms)

Mean amplitudes in the 250–500 ms window were quantified over a centro-parietal ROI (C3/Cz/C4/P3/Pz/P4). A linear mixed-effects model revealed a robust positive effect of sequence position, F(1, 44.39) = 81.45, p < .001, indicating that late responses increased across sequence position. The estimated slope (β = 1.34 µV per position step, 95% CI [0.85, 1.83]) corresponds to an approximate increase of ∼4 µV from the first to the final sequence element. The Domain × Position interaction did not reach significance, F(2, 292) = 2.91, p = .056. Therefore, the primary finding was a robust position-related increase, whereas evidence for domain-specific differences in this increase was weak. A significant main effect of Domain, F(2, 292) = 9.29, p < .001, reflected overall amplitude differences across domains. Figure 3A and 3C illustrate this progressive increase across sequence positions within each domain in central and parietal ROIs.

**Figure 3.**
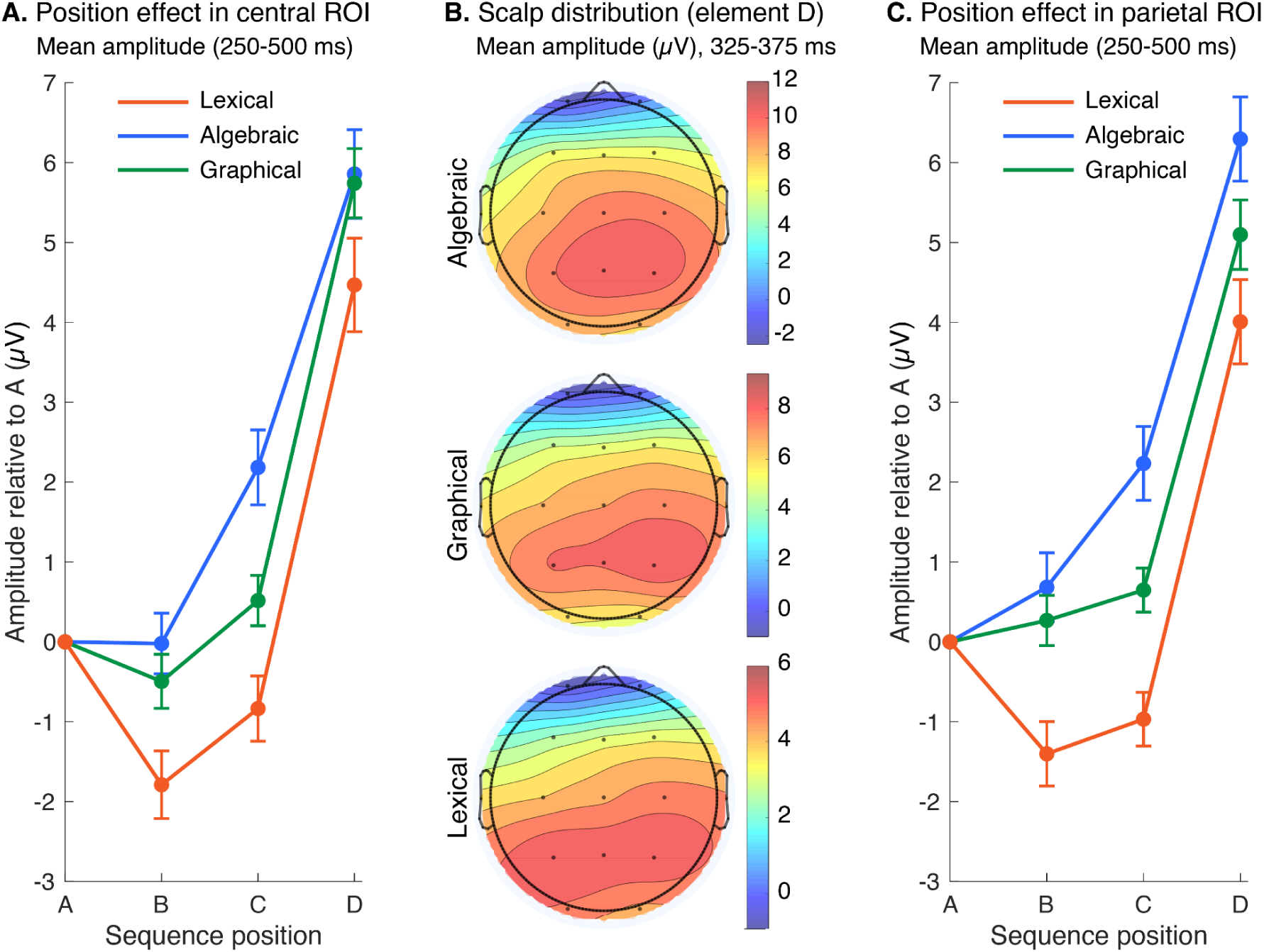
Position Effect and Scalp Distribution of Late-Window Responses Across Regions. Panels A and C illustrate the position effect across representative regional ROIs, complementing the centro-parietal ROI analysis reported in the main text. (A) Mean ERP amplitude in the 250–500 ms window averaged over the central electrodes (C3/Cz/C4), plotted across sequence position (A–D) for the lexical, algebraic, and graphical domains. For visualization, amplitudes are expressed relative to position A within each participant and domain (A = 0). Error bars indicate within-subject SEM (Cousineau-Morey correction). (B) Scalp topographies for the response to the final element (D), showing absolute mean amplitude (i.e., not referenced to A) in a 50-ms window centered on the grand-average peak (∼350 ms; 325–375 ms) for each domain. Topographies are displayed with separate color scales to emphasize the spatial distribution within each domain. (C) Mean ERP amplitude in the 250–500 ms window averaged over the parietal ROI (P3/Pz/P4), plotted across sequence position (A–D) for the lexical, algebraic, and graphical domains. As in panel A, amplitudes are expressed relative to position A within each participant and domain (A = 0).

Peak latency for the final element occurred at approximately 350 ms across domains. Figure 3B shows scalp topographies for the absolute mean amplitude elicited by element D in a 50-ms window centered on this peak (325–375 ms). Across domains, the response showed a broadly centro-parietal distribution consistent with the predefined centro-parietal ROI used in the main analysis.

To further characterize the shape of the position effect within this predefined centro-parietal ROI, a follow-up mixed-effects model compared adjacent increases across the sequence. The effect of Transition was significant, *F*(2, 208) = 81.60, *p* < .001. Bonferroni-corrected pairwise comparisons showed that the C-to-D increase was larger than both the A-to-B increase (mean difference = 4.646 µV, *SE* = 0.385, *p* < .001) and the B-to-C increase (mean difference = 3.708 µV, *SE* = 0.385, *p* < .001). In addition, the B-to-C increase was larger than the A-to-B increase (mean difference = 0.938 µV, *SE* = 0.385, *p* = .047). Thus, within the centro-parietal ROI, the largest increase occurred at the final sequence step. Because element D was also the response-relevant and perceptually marked completion item, this final increase should be interpreted in relation to both contextual completion and task demands.

Given the centro-parietal distribution observed in the topographies and consistent with the literature, we additionally examined whether the position effect varied across scalp regions. A mixed-effects model including ROI (frontal, central, parietal, occipital) revealed a significant ROI × Position interaction, F(3, 1246) = 11.41, p < .001, indicating that the strength of the position effect differed across regions. The Domain × ROI × Position interaction was not significant, F(6, 1246) = 0.40, p = .879, suggesting that this spatial pattern was consistent across domains. Model estimates indicated that the position effect was strongest over central sites (p = .006), remained evident over parietal sites (p = .034), and was weaker over frontal sites (p = .208). Thus, although the largest late amplitudes were observed over parietal electrodes, the graded increase across sequence position was most clearly expressed over central regions.

## Discussion

The present findings indicate that context-sensitive ERP activity across structured sequences was progressive but not uniform. In the 250–500 ms interval, late centro-parietal responses increased across successive elements, with the strongest increase occurring at sequence completion. Although the largest amplitudes were observed over parietal sites, the graded increase across sequence position was most clearly expressed over central regions. Importantly, maximal response amplitude and sensitivity to sequence position did not fully coincide spatially. Because the position-related increase was observed across lexical, algebraic, and graphical sequences, and because evidence for domain-specific differences was limited, the results are compatible with a broadly similar temporal profile across domains.

The present findings extend prior work by characterizing how contextual effects build up across structured sequences in different symbolic domains. Prior work, especially in language, has already shown that responses in this time range vary with graded aspects of contextual support and change as sentence context unfolds across words (Van Petten & Kutas, 1990, 1991; Van Petten & Luka, 2012; Payne et al., 2015; Payne & Federmeier, 2018). More recent accounts likewise emphasize that comprehension unfolds continuously as incoming input is related to an evolving internal representation rather than evaluated only at isolated target positions (Federmeier, 2022). Computational approaches are broadly compatible with this perspective, modeling activity in this range as graded updating as contextual information accumulates (Rabovsky et al., 2018; Michaelov et al., 2024). The present study adds to this literature by characterizing the shape of that buildup within congruent sequences using rigorously controlled four-element sequences with matched structure across domains. At the same time, the present findings do not determine whether the observed activity reflects semantic integration, facilitated lexical access, or a related updating operation, all of which remain debated in the literature (Lau et al., 2009; Rabovsky et al., 2018).

Crucially, the observed pattern did not resemble either a purely uniform accumulation process or a response confined to the final element. Activity increased from A to B to C, indicating that contextual information was incorporated throughout the sequence, in line with earlier work showing that context-sensitive responses can build up gradually as structured input unfolds. This graded buildup was most strongly expressed over central and parietal regions, even though the largest late amplitudes were observed over parietal electrodes. At the same time, the disproportionately large C-to-D transition suggests that sequence completion had a special status. This pattern suggests that contextual integration does not proceed at a constant rate, but instead culminates in a disproportionately strong update at sequence completion.

Work on written sentence comprehension has shown that predictability and plausibility can dissociate not only in the N400 range but also in later positivities, suggesting that sequence-final responses may reflect multiple overlapping constraints rather than a single all-purpose closure mechanism (DeLong et al., 2014). In the present case, one plausible interpretation is that the final element imposed the strongest constraint on the evolving representation, yielding a more fully specified interpretation upon completion of the four-element sequence. On this view, integration does not begin at the final element, but reaches its strongest expression there, consistent with frameworks that link responses in this time range to progressive updating and increasing contextual constraint (Van Petten & Luka, 2012; Rabovsky et al., 2018; Michaelov et al., 2024). The strongest increase at sequence completion should not be interpreted as a trivial consequence of final position alone. Even if the last element involves additional closure-related processing, the progressive increase from A to C indicates that contextual integration was already unfolding across the sequence.

The enhanced response to the final element likely reflects more than a simple motor or response-related process. Because the final element completed the sequence and allowed the preceding context to be evaluated as a coherent whole, it may also have elicited completion– or closure-related processing. Thus, the final-position enhancement is best interpreted as reflecting contextual completion under task-relevant conditions, rather than either a pure integration effect or a purely response-driven effect. More specifically, the final element may have narrowed the set of viable interpretations more strongly than the preceding elements (Stowe et al., 2018; Brothers et al., 2023).

The cross-domain aspect of the findings is consistent with this interpretation. Context-sensitive responses in this latency range have been described most extensively in language, but related effects have also been reported in arithmetic and sequential image processing (Niedeggen et al., 1999; Dickson et al., 2018; Cohn et al., 2012, 2014). The present results extend this idea by showing a comparable temporal profile in algebraic sequences, a symbolic domain that places stronger demands on relational and rule-based processing than the arithmetic expressions more commonly used in prior ERP research. However, these literatures have generally been developed in paradigms that differ in stimulus format, sequence structure, and analytic emphasis, making it difficult to assess directly whether the reported effects reflect a comparable temporal profile of contextual integration. A particular strength of the present study was the use of rigorously controlled four-element sequences that were structurally comparable across domains. This enabled comparisons of how responses unfolded across positions under common constraints. The observation of progressive, non-uniform increases across all three domains therefore suggests that activity in this time range may not be limited to a single stimulus class, but may instead reflect partially shared ERP dynamics associated with the integration of contextual information across structured sequences (Federmeier, 2022; Rabovsky et al., 2018).

The present design is informative because it focuses on how contextual effects accumulate within congruent sequences, rather than relying solely on congruent-incongruent contrasts, which may emphasize violation-related processes more than the graded effects of predictability and contextual support (Lau et al., 2016). This distinction is important because incongruent endings may index failed integration, violation detection, or expectancy mismatch, whereas congruent sequences allow the temporal buildup of a coherent representation to be examined more directly. More broadly, the present approach is consistent with efforts to characterize how neural responses unfold over structured input rather than focusing only on isolated critical events (Alday, 2019). The present pattern is also compatible with the broader idea that centro-parietal activity can exhibit accumulation-like dynamics, although the current design does not permit strong claims about decision-related accumulation in the narrow sense (O’Connell et al., 2012).

The contrast with the early time window further supports this account. Early activity (0–180 ms) showed a consistent posterior-to-anterior gradient across domains, compatible with sensory-perceptual processing, but did not exhibit the progressive position-related scaling observed in the later interval. This dissociation suggests that the buildup observed in the 250–500 ms range was not a simple consequence of low-level stimulus progression, but emerged at a later stage, more closely related to contextual integration. At a broader level, this pattern is consistent with accounts that distinguish modality-specific sensory processing from later N400-range activity linked to context-sensitive meaning construction (Wlotko et al., 2012; Kutas & Federmeier, 2011). From this perspective, the present profile is also compatible with broader accounts that view comprehension as probabilistic event interpretation rather than as a sequence of isolated word-level evaluations (Kuperberg, 2016).

Behavioral performance was consistent with successful task engagement. Accuracy was near ceiling across domains, although reaction times were slower in the lexical domain than in the algebraic and graphical domains. The key neural effect concerned within-sequence scaling across positions, and that scaling was observed in all three domains. This dissociation makes it less likely that the late ERP effect simply reflects global task difficulty or generic response demands. Instead, it is more consistent with the view that the late response varied as contextual information accumulated within the sequence (Van Petten & Kutas, 1990, 1991; Payne & Federmeier, 2018).

Several limitations should be noted. First, the present design used short, highly structured sequences in constrained symbolic domains, so it remains to be determined whether similar temporal profiles would be observed in more naturalistic or temporally extended contexts. Second, the sequences were structurally matched across domains and were designed to provide highly constrained completions within each domain. However, contextual constraint was not quantified using equivalent normative measures for the algebraic and graphical stimuli. Future work should include matched predictability or constraint measures across stimulus classes to better assess domain comparability. Third, although the present findings characterize the temporal shape of contextual buildup, they do not identify the precise computational mechanism responsible for it. The observed profile is consistent with progressive updating and increasing interpretive constraint, but future work combining electrophysiology with formal modeling may help specify the underlying computations more directly. Fourth, the fourth element differed from the preceding elements in several task-relevant respects: it was presented in yellow, remained on the screen for a longer duration, and was the element on which participants made their congruency judgment. These features were introduced to maintain attention, ensure reliable behavioral evaluation, and clearly indicate the response point without adding an additional auditory or visual cue. Nevertheless, these design choices mean that the final-position response cannot be interpreted as a pure effect of contextual integration.

We therefore interpret the enhanced final-position response as reflecting contextual completion under task-relevant conditions, rather than as a pure integration effect. At the same time, because physical stimulus differences are generally expected to exert their strongest influence on early visual components, and because position-related increases were already evident before the final element, the late centro-parietal modulation is unlikely to be explained solely by low-level perceptual salience. However, because color, duration, response relevance, and sequence completion were not independently manipulated, their contributions to the final-position enhancement cannot be fully separated.

Taken together, the present results suggest that contextual integration over structured input is progressive but not uniform. Late responses increased across successive elements, yet the largest increase occurred at sequence completion, indicating that the evolving representation was most strongly constrained at the final step. Although the largest amplitudes were observed over parietal sites, the graded increase across sequence position was also clearly expressed over central regions, suggesting that maximal response magnitude and incremental buildup did not fully overlap spatially. The presence of position-related increases across lexical, algebraic, and graphical sequences, with limited evidence for domain-specific differences, is consistent with the view that centro-parietal ERP activity in the 250–500 ms window tracks contextual buildup during structured sequence processing.

## Conclusion

The present findings show that contextual integration across structured sequences followed a progressive but non-uniform temporal profile. In the 250–500 ms interval, late responses increased across successive elements, with the strongest increase occurring when the sequence reached its response-relevant completion point. Although the largest amplitudes were observed over parietal sites, the graded increase across sequence position was most clearly expressed over central regions. Importantly, maximal response amplitude and sensitivity to sequence position did not fully coincide spatially. This pattern was observed across lexical, algebraic, and graphical domains, with no strong evidence for domain-specific differences in the position-related increase, suggesting that context-sensitive activity in this time range may reflect partially shared electrophysiological dynamics associated with the progressive integration of contextual information across structured symbolic sequences. This pattern contrasts with earlier sensory activity, which did not show comparable position-related scaling. These findings highlight the importance of characterizing not only whether contextual effects are present, but also how they unfold across structured sequential input.

## Declarations

### Funding

This research did not receive any specific grant from funding agencies in the public, commercial, or not-for-profit sectors.

### Competing interests

The authors declare no competing interests.

### Ethics approval

The study was conducted in accordance with the Declaration of Helsinki and was approved by the Ethics Committee of the Institute of Neuroscience, University of Guadalajara.

### Consent to participate

All participants provided written informed consent.

### Consent for publication

Not applicable.

### Data, materials, and code availability

De-identified preprocessed ERP data, stimulus materials, and analysis code will be made publicly available upon publication in an open repository.

Analysis code and materials will also be available at: https://github.com/LupitaYanez/symbolic_domains.

## Author Contributions

María Guadalupe Yáñez-Ramos: Conceptualization, Methodology, Formal analysis, Investigation, Data curation, Writing – original draft, Visualization.

Daniel Zarabozo Enríquez de Rivera: Conceptualization, Supervision, Methodology, Writing – review & editing.

Andrés Antonio González Garrido: Supervision, Writing – review & editing.

## Notes

https://github.com/LupitaYanez/symbolic_domains

### Competing Interest Statement

The authors have declared no competing interest.

### Summary of Updates

This version has been revised to clarify the interpretation of the ERP effect in the 250-500 ms window, emphasizing that the analysis examines context-sensitive activity within congruent sequences rather than a conventional N400 incongruity effect. The title and abstract were updated to better reflect the progressive nature of the position-related response. The Introduction and Discussion were revised to clarify the rationale for focusing on congruent trials and to distinguish contextual buildup from congruent-incongruent contrasts. The Results and Discussion were also revised to provide a more cautious interpretation of domain-related differences and of the enhanced response to the final, response-relevant completion item. Additional methodological clarification was added regarding the color marking and longer presentation duration of the final element. Minor wording changes were made throughout to improve clarity.

## References

1. Annett, M. (1970). A classification of hand preference by association analysis. British Journal of Psychology, 61(3), 303–321. 10.1111/j.2044-8295.1970.tb01248.x

2. Alday, P. M. (2019). M/EEG analysis of naturalistic stories: A review from speech to language processing. Language, Cognition and Neuroscience, 34(4), 457–473. 10.1080/23273798.2018.1546882

3. Brothers, T., Morgan, E., Yacovone, A., & Kuperberg, G. R. (2023). Multiple predictions during language comprehension: Friends, foes, or indifferent companions? Cognition, 241, Article 105602. 10.1016/j.cognition.2023.105602

4. Brothers, T., Swaab, T. Y., & Traxler, M. J. (2015). Effects of prediction and contextual support on lexical processing: Prediction takes precedence. Cognition, 136, 135–149. 10.1016/j.cognition.2014.10.017

5. Cohn, N., Jackendoff, R., Holcomb, P. J., & Kuperberg, G. R. (2014). The grammar of visual narrative: Neural evidence for constituent structure in sequential image comprehension. Neuropsychologia, 64, 63–70. 10.1016/j.neuropsychologia.2014.09.018

6. Cohn, N., Paczynski, M., Jackendoff, R., Holcomb, P. J., & Kuperberg, G. R. (2012). Structure and meaning in sequential image comprehension. Cognitive Psychology, 65(3), 229–272. 10.1016/j.cogpsych.2012.06.003

7. Dambacher, M., Kliegl, R., Hofmann, M., & Jacobs, A. M. (2006). Frequency and predictability effects on event-related potentials during reading. Neuroscience Letters, 402(1–2), 89–93. 10.1016/j.neulet.2006.03.051

8. DeLong, K. A., Quante, L., & Kutas, M. (2014). Predictability, plausibility, and two late ERP positivities during written sentence comprehension. Neuropsychologia, 61, 150–162. 10.1016/j.neuropsychologia.2014.06.016

9. Delorme, A., & Makeig, S. (2004). EEGLAB: An open-source toolbox for analysis of single-trial EEG dynamics including independent component analysis. Journal of Neuroscience Methods, 134(1), 9–21. 10.1016/j.jneumeth.2003.10.009

10. Dickson, D. S., Federmeier, K. D., & Kutas, M. (2018). When “2 × 4” is meaningful: The N400 and P300 reveal operand format effects in multiplication verification. Psychophysiology, 55(12), Article e13212. 10.1111/psyp.13212

11. Federmeier, K. D. (2022). Connecting and considering: Electrophysiology provides insights into comprehension. Psychophysiology, 59(1), Article e13940. 10.1111/psyp.13940

12. Knoeferle, P., Urbach, T. P., & Kutas, M. (2011). Comprehending how visual context influences incremental sentence processing: Insights from ERPs and picture-sentence verification. Psychophysiology, 48(4), 495–506. 10.1111/j.1469-8986.2010.01080.x

13. Kuperberg, G. R. (2016). Separate streams or probabilistic inference? What the N400 can tell us about the comprehension of events. Language, Cognition and Neuroscience, 31(5), 602–616. 10.1080/23273798.2015.1130233

14. Kutas, M., & Federmeier, K. D. (2011). Thirty years and counting: Finding meaning in the N400 component of the event-related brain potential (ERP). Annual Review of Psychology, 62, 621–647. 10.1146/annurev.psych.093008.131123

15. Kutas, M., & Hillyard, S. A. (1980). Reading senseless sentences: Brain potentials reflect semantic incongruity. Science, 207(4427), 203–205. 10.1126/science.7350657

16. Lau, E. F., Almeida, D., Hines, P. C., & Poeppel, D. (2009). A lexical basis for N400 context effects: Evidence from MEG. Brain and Language, 111(3), 161–172. 10.1016/j.bandl.2009.08.007

17. Lau, E. F., Namyst, A., Fogel, A., & Delgado, T. (2016). A direct comparison of N400 effects of predictability and incongruity in adjective-noun combination. Collabra, 2(1), Article 13. 10.1525/collabra.40

18. Lopez-Calderon, J., & Luck, S. J. (2014). ERPLAB: An open-source toolbox for the analysis of event-related potentials. Frontiers in Human Neuroscience, 8, Article 213. 10.3389/fnhum.2014.00213

19. Luck, S. J. (2014). An introduction to the event-related potential technique (2nd ed.). MIT Press.

20. Michaelov, J. A., Bardolph, M. D., Bergen, B. K., & Coulson, S. (2024). Language model surprisal explains multiple N400 effects. Neurobiology of Language, 5(1), 107–136. 10.1162/nol_a_00137

21. Niedeggen, M., Rösler, F., & Jost, K. (1999). Processing of incongruous mental calculation problems: Evidence for an arithmetic N400 effect. Psychophysiology, 36(3), 307–324. 10.1017/S0048577299981184

22. Nour Eddine, S., Brothers, T., Wang, L., Spratling, M., & Kuperberg, G. R. (2024). A predictive coding model of the N400. Cognition, 246, Article 105755. 10.1016/j.cognition.2024.105755

23. O’Connell, R. G., Dockree, P. M., & Kelly, S. P. (2012). A supramodal accumulation-to-bound signal that determines perceptual decisions in humans. Nature Neuroscience, 15(12), 1729–1735. 10.1038/nn.3248

24. Payne, B. R., & Federmeier, K. D. (2018). Contextual constraints on lexico-semantic processing in aging: Evidence from single-word event-related brain potentials. Neuroscience Letters, 685, 117–122. 10.1016/j.neulet.2018.08.040

25. Payne, B. R., Lee, C.-L., & Federmeier, K. D. (2015). Revisiting the incremental effects of context on word processing: Evidence from single-word event-related brain potentials. Psychophysiology, 52(11), 1456–1469. 10.1111/psyp.12515

26. Peirce, J. W. (2009). Generating stimuli for neuroscience using PsychoPy. Frontiers in Neuroinformatics, 2, Article 10. 10.3389/neuro.11.010.2008

27. Rabovsky, M., Hansen, S. S., & McClelland, J. L. (2018). Modelling the N400 brain potential as change in a probabilistic representation of meaning. Nature Human Behaviour, 2(9), 693–705. 10.1038/s41562-018-0406-4

28. Rokszin, A. A., Győri-Dani, D., Nyúl, L. G., & Csifcsák, G. (2016). Electrophysiological correlates of top-down effects facilitating natural image categorization are disrupted by the attenuation of low spatial frequency information. International Journal of Psychophysiology, 100, 19–27. 10.1016/j.ijpsycho.2015.12.006

29. Stowe, L. A., Kaan, E., Sabourin, L., & Taylor, R. C. (2018). The sentence wrap-up dogma. Cognition, 176, 232–247. 10.1016/j.cognition.2018.03.011

30. Tanner, D., Morgan-Short, K., & Luck, S. J. (2015). How inappropriate high-pass filters can produce artifactual effects and incorrect conclusions in ERP studies of language and cognition. Psychophysiology, 52(8), 997–1009. 10.1111/psyp.12437

31. Van Petten, C., & Kutas, M. (1990). Interactions between sentence context and word frequency in event-related brain potentials. Memory & Cognition, 18(4), 380–393. 10.3758/BF03197127

32. Van Petten, C., & Kutas, M. (1991). Influences of semantic and syntactic context on open– and closed-class words. Memory & Cognition, 19(1), 95–112. 10.3758/BF03198496

33. Van Petten, C., & Luka, B. J. (2012). Prediction during language comprehension: Benefits, costs, and ERP components. International Journal of Psychophysiology, 83(2), 176–190. 10.1016/j.ijpsycho.2011.09.015

34. Wlotko, E. W., Federmeier, K. D., & Kutas, M. (2012). So that’s what you meant! Event-related potentials reveal multiple aspects of context use during construction of message-level meaning. NeuroImage, 62(1), 356–366. 10.1016/j.neuroimage.2012.04.054

35. Yáñez-Ramos, M. G., Zarabozo, D., & Varela, J. (2022). Validación de oraciones con cierres congruentes e incongruentes en población mexicana. Investigación y Ciencia de la Universidad Autónoma de Aguascalientes, 86. 10.33064/iycuaa2022863534

